# Graph neural networks learn emergent tissue properties from spatial molecular profiles

**DOI:** 10.1101/2022.12.08.519537

**Authors:** David S. Fischer, Mayar Ali, Sabrina Richter, Ali Ertürk, Fabian Theis

**Affiliations:** Institute of Computational Biology, Helmholtz Zentrum München, 85764 Neuherberg, Germany; TUM School of Life Sciences Weihenstephan, Technical University of Munich, 85354 Freising, Germany; Institute for Tissue Engineering and Regenerative Medicine, Helmholtz Zentrum München, 85764 Neuherberg, Germany; Graduate School of Systemic Neurosciences, Ludwig Maximilian University of Munich, 80539 Munich, Germany; Department of Mathematics, Technical University of Munich, 85748 Garching bei München, Germany

**Author notes:** These authors contributed equally (David S. Fischer, Mayar Ali, Sabrina Richter).

## Abstract

Tissue phenotypes such as metabolic states, inflammation, and tumor properties are functions of molecular states of cells that constitute the tissue. Recent spatial molecular profiling assays measure tissue architecture motifs in a molecular and often unbiased way and thus can explain some aspects of emergence of these phenotypes. Here, we characterize the ability of graph neural networks to model tissue-level emergent phenotypes based on spatial data by evaluating phenotype prediction across model complexities. First, we show that immune cell dispersion in colorectal tumors, which is known to be predictive of disease outcome, can be captured by graph neural networks. Second, we show that breast cancer tumor classes can be predicted from gene expression alone without spatial information and are thus too simplistic a phenotype to require a complex model of emergence. Third, we show that representation learning approaches for spatial graphs of molecular profiles are limited by overfitting in the prevalent regime of up to 100s of images per study. We address overfitting with within-graph self-supervision and illustrate its promise for tissue representation learning as a constraint for node representations.

## Introduction

The high molecular resolution provided by single-cell RNA-seq (scRNA-seq) has put the cell as a functional unit in the focus of recent advances in tissue biology^1^. However, interactions between cells and emergent properties of the tissue beyond the length scale of a cell are largely lost in assays that are based on dissociated tissues. Spatial molecular profiling assays with single-cell resolution can fill this gap between cell and tissue biology by providing descriptions of tissues as graphs of cells^2^. For example, emergent properties of cells in tissue niches can be modeled in such spatial graphs^3^. Sample-level labels often reflect complex tissue phenotypes, such as metabolic properties or disease states. In the absence of spatial information, a supervised model on sample-level labels can detect cell states and frequencies thereof that correlate with the tissue phenotype. Spatial information and graphs of cells extend the capability of such supervised models to not only detect cellular phenotypes that correlate with the labels, but also motifs of tissue architecture, such as cellular niches^4^. Because of their explicit representation of cells as constituent building blocks of a tissue, graph neural networks promise to be more interpretable with respect to niches than convolutional neural networks on tissue images and, indeed, have been recently successfully deployed for tumor phenotype prediction from spatial proteomics data^5–7^. Here, we perform a comparative ablation study over spatial features and single-cell resolution for graph neural networks that predict tumor phenotypes from spatial graphs with gene expression or categorical cell type node features. We use this analysis to show that spatial motives of cell types that are predictive of tumor labels are encoded in cell-wise gene expression, thus delineating whether spatial edges need to be modeled.

## Results

### Graph neural networks model tissue phenotypes

We introduce graph neural networks over spatial graphs of cells with an aggregation function across node states to model graph-level labels that represent tissue phenotypes (Fig. 1a). The input node features can either be gene expression vectors per cell or coarsened gene expression features, such as categorical cell type labels (Fig. 1b). This model class captures both molecular information of individual cells as input node feature vectors and the architecture of the tissue through the spatial proximity graph. Previously, such graph neural network architectures have been used to model tumor grades based on H&E images^8^ and spatial proteomics^5–7^. We consider examples of graph-level supervision on three data sets: Two large cohorts of imaging mass cytometry (IMC) of breast cancer biopsy data stratified by tumor grade and other tumor characteristics (*IMC - Jackson*^*9*^ with 559 images from 350 patients and *IMC - METABRIC*^*9*^ with 500 images from 454 patients) and a cohort of CODEX samples of colorectal cancer biopsies stratified by tumor classification and disease outcome (*CODEX - colorectal cancer*^*10*^, 140 images from 35 patients). We focussed on tumor grade prediction on the *IMC - Jackson* and *IMC - METABRIC* datasets. In these published breast cancer data sets, the grade label was previously assigned by a pathologist to the tumors based on standard histology and the tumor section assayed with IMC were chosen to represent this decision. In both cases, we considered the problem of distinguishing grade 1, 2 and 3 tumors without introducing class weighting. We considered the prediction of binary tumor classes defined based on the presence of diffuse inflammatory infiltration in the *CODEX - colorectal cancer* data set (Fig. 1c). In all data sets, we performed training, validation and test data splits by patients and grouped all images of a patient into one partition. We chose neighborhood sizes for spatial graphs (resolution) based on the average node degree distribution^3^ (Supp. Fig. 1). We standardized all features globally across all data as we found this to best reflect cell type and condition information (Supp. Fig. 2).

**Figure 1:**
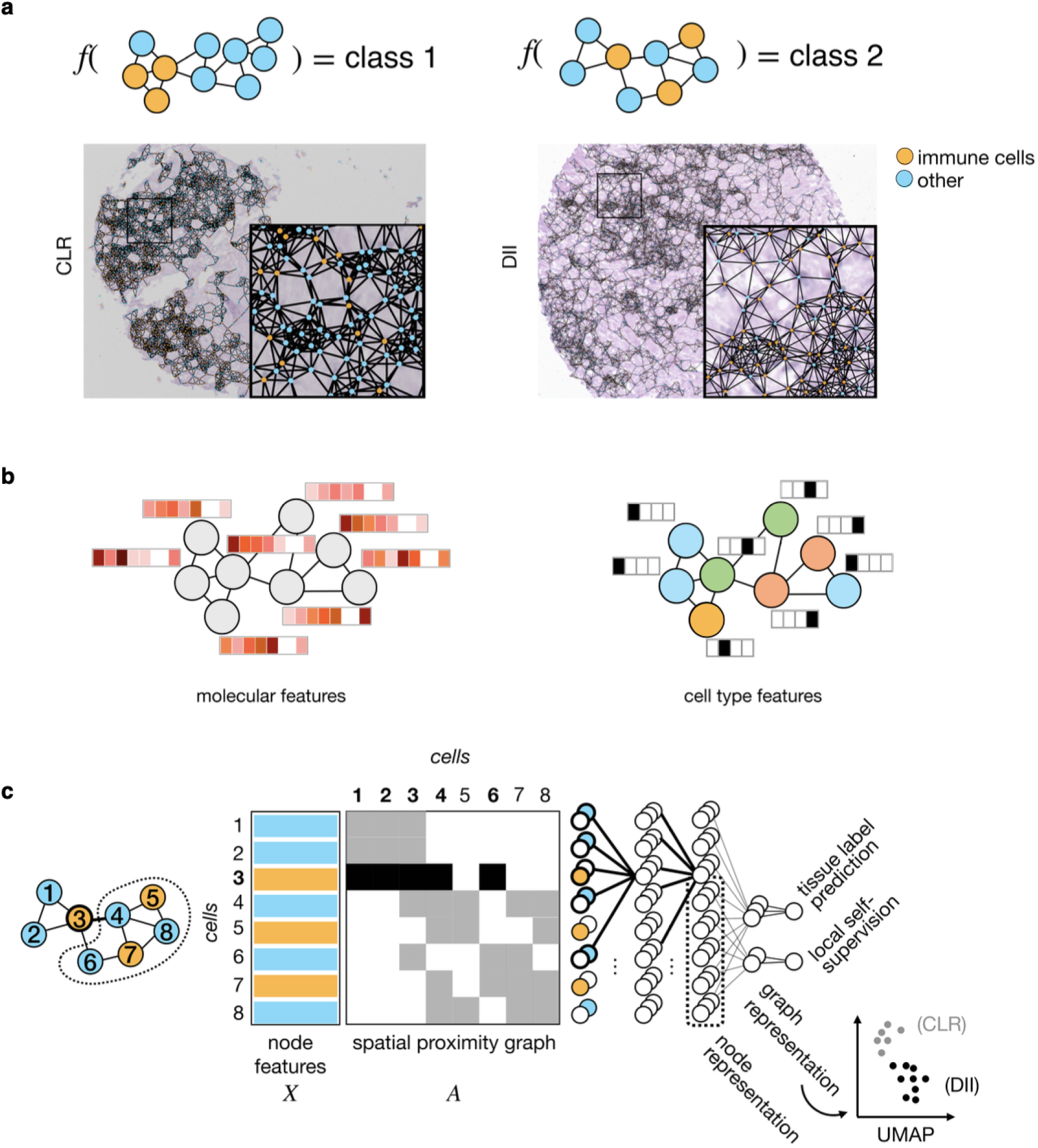
A graph neural networks model of tissue phenotypes. **(a)** Tissue-level phenotypes are functions of the architecture of the tissue. In this case, two colorectal tumor classes, Crohn’s-like reaction (CLR) and diffuse inflammatory infiltration (DII), can be distinguished based on the spatial distribution of immune cells. This tumor label cannot be inferred based on frequencies of cell types that would be available in dissociation-based protocols, but only based on the spatial distribution of cells^10^. One example image from the *CODEX - colorectal cancer* dataset for each class. **(b)** A graphical representation of a tissue graph with different node features, molecular gene expression profiles (left) and one-hot encoding of node cell type (right). **(c)** The spatial context of each cell can be formally represented by a graph in which edges are weighted based on the distance between nodes. Each sample can be represented as one such graph, where nodes are colored by the measured cell features. We perform prediction with a model that consists of graph neural network layers to produce node embeddings, followed by pooling over nodes and a final classification network. In addition, the node embeddings of connected components of nodes on the spatial proximity graph can be aggregated for local self-supervision tasks, such as reconstruction of adjacent clusters’ cell type composition. *dotted line*: connected component of nodes on spatial proximity graph. The spatially-aware graph embedding can be visualized with a UMAP in which each point reflects one graph (image) and depicts separation of samples by the tumor class (CLR and DII) here.

### Predictive features of tumor architecture are defined by an ablation study

When applying the GNN models to the three tumor settings described above, we found graph neural networks with node state aggregation to be predictive of the considered tissue labels (Fig. 2). We performed an ablation study over single-cell and spatial information for a tissue phenotype to investigate if these models require the spatial proximity graph, thus testing if they capture emergent properties of tissue architecture. We designed the ablation based on two baseline experiments, omitting spatial structure and cell resolution from spatial graphs of single cells (Fig. 2b). The first baseline model uses pseudo bulk data, which consists of mean gene expression vectors across an entire image. We used fully connected dense neural networks including linear models (MLP) to predict tissue-level phenotypes on pseudo-bulk data (Online Methods). The second baseline model uses *in silico* dissociated single-cell data, which corresponds to removing the observed cellular gene expression vectors from their spatial context to yield independent observations. We modeled this *in silico* dissociated single-cell data with a multi-instance (MI) model that aggregate cell-wise embeddings (Online Methods). We compared the performance of these baseline models with spatial graph convolutional networks (GCN) that leverages spatial information (Fig. 1c). We considered GCNs trained with cross entropy loss functions on the tumor phenotype. In addition, we trained GCNs with an additional self-supervision task (GCN-SS). This self-supervision task consists of predicting cell type composition in adjacent regions in the graph from local graph embeddings, in analogy to predicting held-out patches in self-supervision in images (Online Methods).

**Figure 2:**
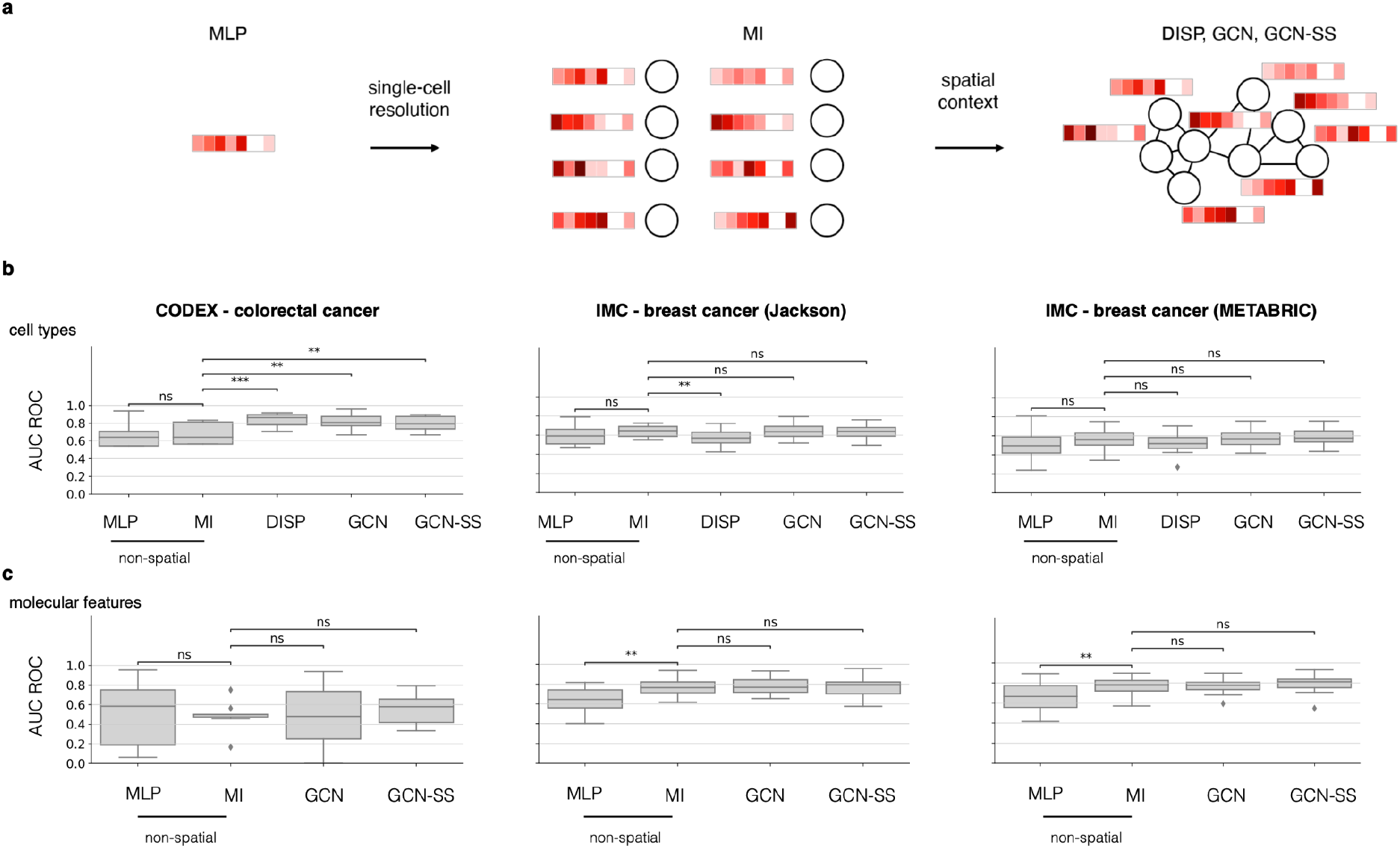
Graph neural networks predict whole-slide tumor phenotypes from cellular architecture. **(a)** Design of the ablation study. MLP models only have access to the average node feature vector of the graph. MI models have access to single-cell-resolved but *in silico* dissociated data from the observed spatial graph. Spatially aware models, DISP, GCN, and GCN-SS, have access to node features and the spatial proximity graph. *MLP*: Multi-layer perceptron based on graph-wide summary statistics on features. *MI*: Multi-instance model on nodes of graph. *DISP*: Dispersion multi-layer perceptron based on average graph niches features. *GCN*: Graph convolutional network. *GCN-SS*: Graph convolutional network with additional self-supervision loss. **(b, c)** Three separate applications of graph neural networks to predict tumor phenotypes on the IMC - breast cancer (Jackson), IMC - breast cancer (METABRIC) and CODEX - colorectal cancer datasets. Model complexity ablation study on classification performance on breast cancer grade prediction and colorectal cancer group prediction using two feature spaces, molecular features and cell types. Shown is the area-under-curve of the receiver-operator characteristic curve (AUC ROC) using the binary cell type features (immune and non-immune) **(b)** and molecular profile features **(c)** across six-fold cross-validation for the best performing hyper-parameter set for each model class for best models selected on train loss.

First, we considered one-hot encoded cell type labels as input node features. MI models did not outperform pseudo-bulk models on coarse binary cell type labels (immune and non-immune cells, Fig. 2b) and on more fine-grained author-supplied labels (Supp. Fig. 3a-d). It was previously reported that the spatial distribution of immune cells in colorectal cancer is predictive of disease outcome and is used to stratify tumors^5,10^. Indeed, we found that an MLP trained on an immune cell dispersion estimate per image was more predictive of tumor class in test images of colorectal cancer than the baseline pseudo-bulk model that only reflects composition but not architecture (Fig. 2c, Online Methods). This performance was matched by GCN and GCN-SS (Fig. 2c), demonstrating that these models can independently detect this tissue architecture feature. In contrast, the spatial distribution of immune cells in breast cancer was not predictive of tumor class (Fig. 2c), potentially reflecting differences in tumor biology and measurement characteristics.

Cell type labels are a coarsening of cell-wise gene expression measurements. Within-cell type gene expression variation is in parts explained by spatial niches^3^. We considered gene expression as input feature vectors to ablate over spatial features in the absence of confounding by cell type classification choices. First, we found that MI models outperform pseudo-bulk models on test images of the *IMC - Jackson* dataset significantly (average accuracy difference 0.105), not significantly on the *IMC - METABRIC* and *CODEX - colorectal cancer* datasets (average accuracy difference 0.081 and 0.094, respectively). This performance of MI models indicates that information on tissue-level phenotypes is captured by single-cell-resolved data (Fig. 2c). Second, we found that GCN models do not outperform MI models in the same setting (Fig. 2c). This result shows that predictors of the considered tissue phenotypes are sufficiently captured by non-spatial single-cell gene expression states. One interpretation of this result is that the predictive tissue architecture motives are already encoded in cell-wise gene expression vectors, which would be consistent with the previously demonstrated ability of spatial graphs to explain parts of within-cell-type gene expression variation^3^. Indeed, MI models were significantly better when trained on gene expression vectors as opposed to one-hot encoded cell type node representations on the *IMC - Jackson* data and *IMC - METABRIC* data (Supp. Fig. 3e), demonstrating that this information inherent single-cell resolution is lost upon coarse graining gene expression to cell types.

We noticed that the test performance of the optimal hyperparameter set depended on whether we selected models based on training data loss or validation data loss and deteriorated if hyperparameters were chosen on validation data (Supp. Fig. 4d). The two breast cancer data sets were big enough to include a validation data set. We used 10% of the total data as validation data. The small validation dataset size of a few held-out images may result in stochastic effects in model selection that are detrimental to finding good hyperparameters and we continued to investigate models selected on train data performance. We noticed that the training loss plateaued early in many model fits (Supp. Fig. 5) and hypothesized that GCNs overfit patients during training because of the low number of total samples and the large content of information in the input data. We added further sample-level labels in a multi-task set-up to the GCNs trained on the *IMC - breast cancer* and *IMC - METABRIC* data sets but did not find this to improve the prediction accuracy on test data (Supp. Fig. 6).

### Sample embeddings reflect tumor phenotypes

We sought to interpret the learnt tissue representations of GCNs through sample embeddings (Fig. 1c). After node-pooling, graph-based neural networks yield spatially-aware embeddings of an entire graph that can be used to compare different samples. Interestingly, the graph-embeddings show a continuous manifold of tumor grades ranging from grade 1 via grade 2 to grade 3 in the *IMC - Jackson* and *IMC - METABRIC* datasets (Fig. 3a,b). Note that the order of tumor grades is not encoded in the categorical multi-class prediction problem and was learned correctly by the model, with grade 2 lying between grade 1 and 3 (Fig. 3a,b). The graph-embeddings of grade 1 and 2 tend to overlap in the *IMC - METABRIC* dataset which reflects their anatomic similarity (Fig. 3b). The sample embedding separates both tumor classes in the *CODEX - colorectal cancer* data sets (Fig. 3c).

**Figure 3:**
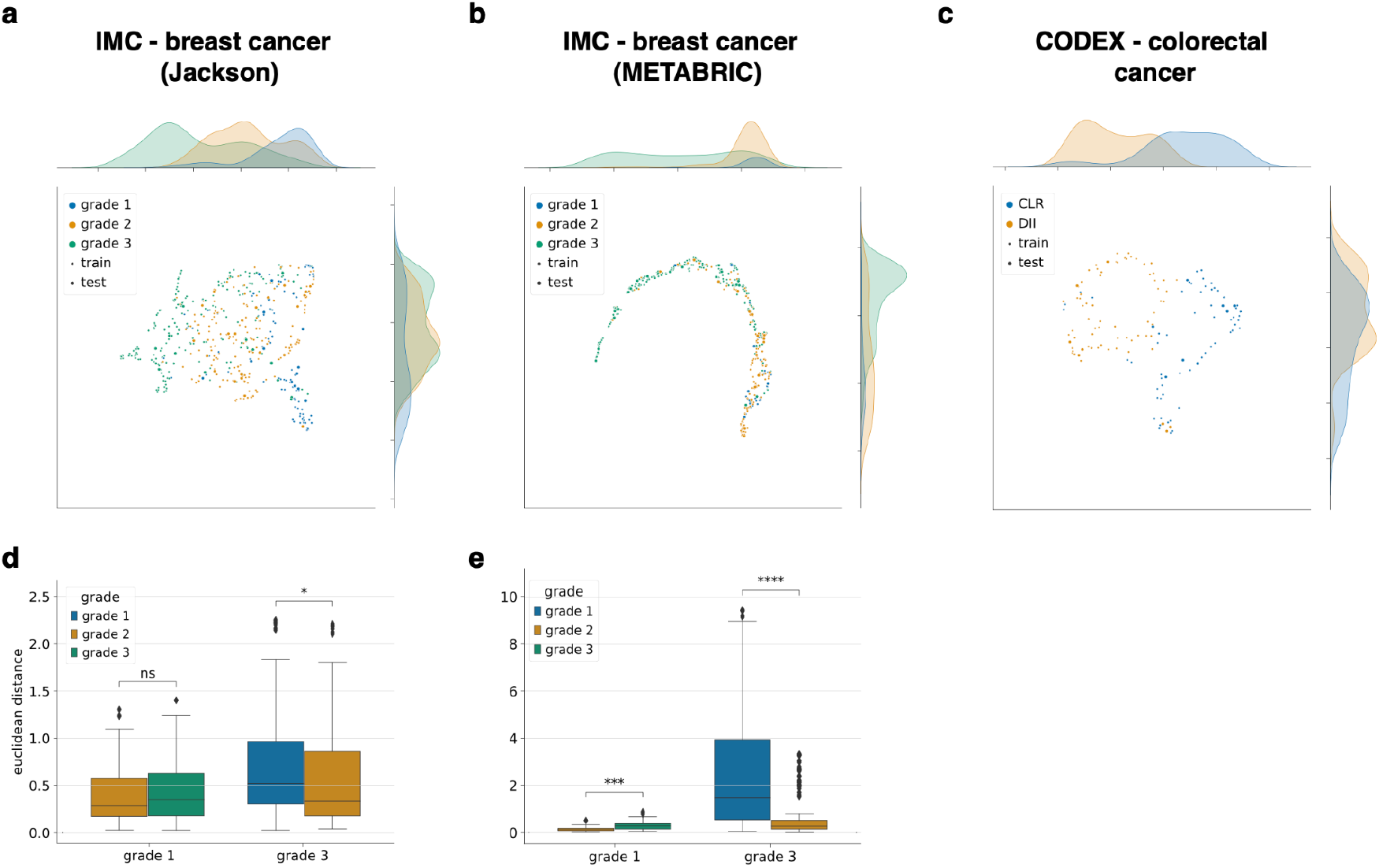
Graph neural networks interpolate tumor states. UMAP of graph embedding of GCN-SS of training and test data with class label superimposed for **(a)** IMC - breast cancer (Jackson), **(b)** IMC - breast cancer (METABRIC) and **(c)** CODEX - colorectal cancer datasets. Each point is one graph. **(d, e)** The euclidean distances between the different class embeddings of the **(d)** IMC - breast cancer (Jackson) and **(e)** IMC - breast cancer (METABRIC).

### Self-supervision mitigates overfitting on the node level

Not having found anomalies on the level of sample embeddings that could explain overfitting, we next considered node embeddings. We considered the distribution of cell types and images of each node in a UMAP fit to the embedding of 52 test images in both the input layer and the final node embedding layer of a GCN and a GCN-SS (Fig. 4). Indeed, we found traces of overfitting to images on the level of node embeddings as a strong separation of nodes of each image in the last node embedding layer of the GCN. We found that the self-supervision loss constrains node embeddings by increasing integration of node embeddings across images (Fig. 4a,b). We quantified this domain correction through self-supervision using data integration metrics from scRNA-seq analysis^11^. We found image integration to improve as measured by data integration metrics^11^ to decrease when adding a self-supervision loss in a GCN to a graph-level label loss (Fig. 4c, Supp. Fig. 8). Overfitting was less prominent for MI models when compared to GCNs as evaluated by data integration metrics (Supp. Fig. 8), demonstrating that the expressiveness of GCNs poses a challenge here. In summary, we showed that self-supervision in graph representation learning is a powerful mechanism to constrain node embeddings and a promising avenue to improve predictive performance in the future.

**Figure 4:**
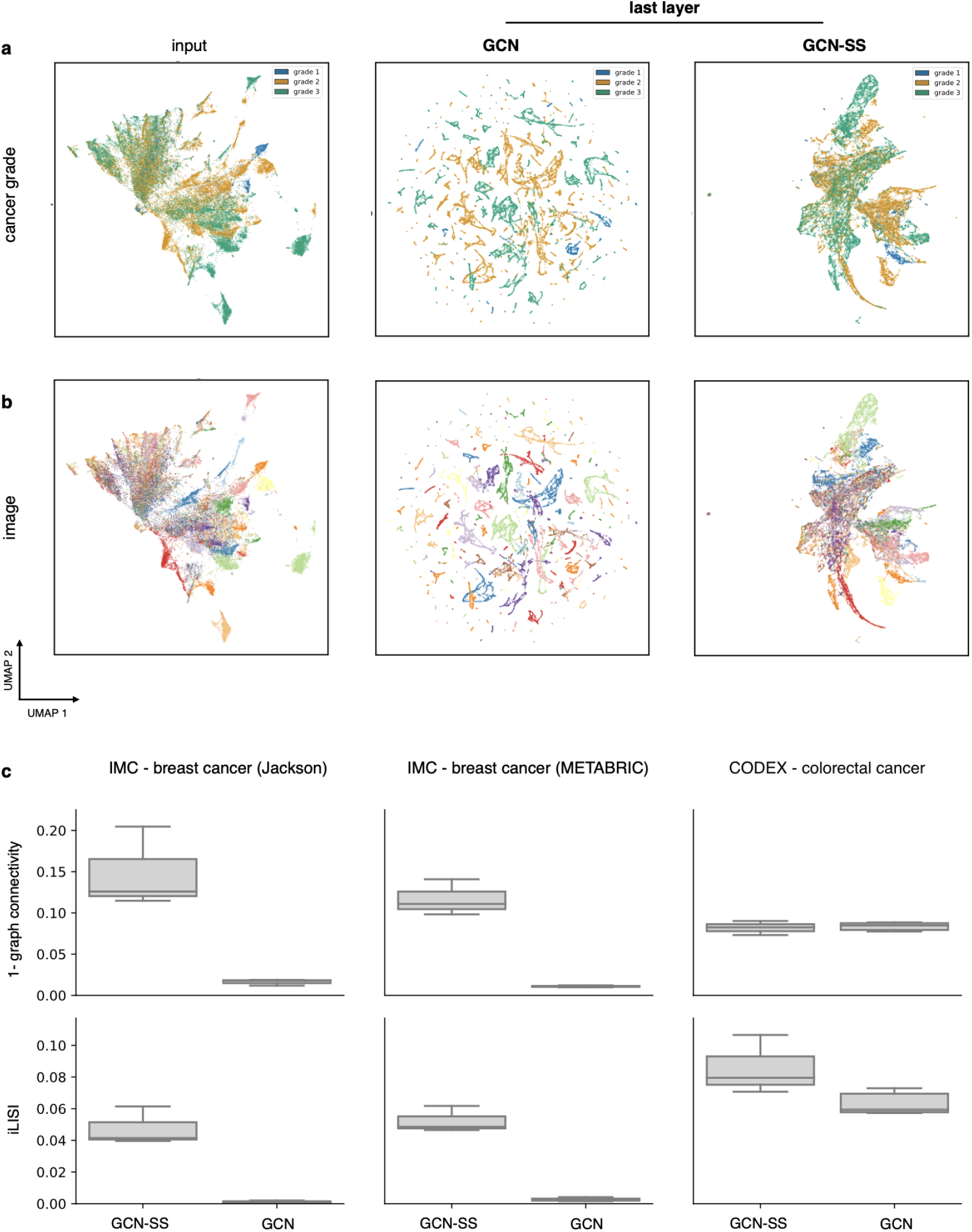
Auxiliary self-supervision loss stabilizes node embeddings. **(a**,**b)** UMAPs of node embeddings by layer: input, last GCN layer in a GCN model and last GCN layer in a GCN-SS with two GCN layers trained on the *IMC - breast cancer* data *(Jackson)*. Superimposed are **(a)** the cancer grade of the corresponding graph and **(b)** the image (graph) to which the node belongs. **(c)** Focusing only on GCN and GCN-SS here, we show the *iLISI and graph connectivity* integration metrics on the three datasets (N=3 cross validations). For *iLISI* and *1 - graph connectivity*, higher values imply better integration, while low values result from distribution differences in the considered batches.

## Discussion

We benchmarked the ability of graph neural networks to predict tissue-level phenotypes based on single-cell-resolved spatial molecular profiling data sets. We detected spatial patterns of immune cells in colorectal cancer related to tumor class with graph neural networks. In breast cancer, the spatial information was sufficiently well represented in the cell-wise gene expression observations so that the predictive performance of GCNs is matched by non-spatial baseline models. This may be, in part, due to the simplicity of the currently available tissue phenotype labels and to the small number of images per study that limits the complexity of representations learned. We discussed self-supervision as a means to improve graph-level representation learning in this regime of low number of data points where overfitting is hard to avoid. We validated our findings in two comparable breast cancer cohorts and a colorectal cancer cohort, using tumor phenotypes as sample labels. Different tissue phenotypes may provide harder supervision problems and thus may give rise to a stronger relative performance of GCNs with respect to baseline models. Metabolic tissue properties and tissue response to perturbation may provide for such complex physiological read-outs.

Beyond the choice of tissue phenotype, there are many model architecture design choices to be considered in the future: First, we considered GCN kernels here. Graph attention^12^ and other graph kernels with more degrees of freedom may be more sensitive to complex tissue niche motives. Second, pooling across the graph may be performed globally or hierarchically, as previously also discussed by Wu *et al*.^*5*^. Aggregation of information across a graph becomes even more relevant if larger graphs are considered, which may for example become available in the context of tissue clearing^13^. Integration of spatial profiling data with deep molecular profiles from scRNA-seq data may provide for higher resolution in the gene expression input feature space to GCNs in the future^14^.

## Online methods Data

### IMC - breast cancer (Jackson)

The breast cancer dataset (Jackson *et al*.^*9,15*^ with 559 images from 350 patients) was measured with IMC. The dataset consists of samples from three breast cancer grades, grade 1 (114 images), grade 2 (214 images) and grade 3 (231 images). Here, 34 proteins in a panel specific to breast cancer microenvironment were simultaneously measured. We used the segmentation provided by Jackson *et al*.. We used the following channels: 1021522Tm169Di EGFR, 1031747Er167Di ECadhe, 112475Gd156Di Estroge, 117792Dy163Di GATA3, 1261726In113Di Histone, 1441101Er168Di Ki67, 174864Nd148Di SMA, 1921755Sm149Di Vimenti, 198883Yb176Di cleaved, 201487Eu151Di cerbB, 207736Tb159Di p53, 234832Lu175Di panCyto, 3111576Nd143Di Cytoker, Nd145Di Twist, 312878Gd158Di Progest, 322787Nd150Di cMyc, 3281668Nd142Di Fibrone, 346876Sm147Di Keratin, 3521227Gd155Di Slug, 361077Dy164Di CD20, 378871Yb172Di vWF, 473968La139Di Histone, 651779Pr141Di Cytoker, 6967Gd160Di CD44, 71790Dy162Di CD45, 77877Nd146Di CD68, 8001752Sm152Di CD3epsi, 92964Er166Di Carboni, 971099Nd144Di Cytoker, 98922Yb174Di Cytoker, phospho Histone, phospho S6, phospho mTOR, Area. Jackon *et al*. annotated the following cell types: B cells, T and B cells, T cells, macrophages, T cells, macrophages, endothelial, vimentin hi stromal cell, small circular stromal cell, small elongated stromal cell, fibronectin hi stromal cell, large elongated stromal cell, SMA hi vimentin hi stromal cell, hypoxic tumor cell, apoptotic tumor cell, proliferative tumor cell, p53+ EGFR+ tumor cell, basal CK tumor cell, CK7+ CK hi cadherin hi tumor cell, CK7+ CK+ tumor cell, epithelial low tumor cell, CK low HR low tumor cell, CK+ HR hi tumor cell, CK+ HR+ tumor cell, CK+ HR low tumor cell, CK low HR hi p53+ tumor cell and myoepithelial tumor cell. We coarsened the cell types into B cells, T and B cells, T cells, macrophages, T cells, macrophages, endothelial, stromal cells (vimentin hi stromal cell, small circular stromal cell, small elongated stromal cell, fibronectin hi stromal cell, large elongated stromal cell, SMA hi vimentin hi stromal cell) and tumor cells (hypoxic tumor cell, apoptotic tumor cell, proliferative tumor cell, p53+ EGFR+ tumor cell, basal CK tumor cell, CK7+ CK hi cadherin hi tumor cell, CK7+ CK+ tumor cell, epithelial low tumor cell, CK low HR low tumor cell, CK+ HR hi tumor cell, CK+ HR+ tumor cell, CK+ HR low tumor cell, CK low HR hi p53+ tumor cell, myoepithelial tumor cell).

### IMC - breast cancer (METABRIC)

The breast cancer METABRIC cohort (Ali et al.^15^ with 500 images from 467 patients) was collected with IMC. Here, 37 proteins in formalin-fixed, paraffin-embedded breast tumor samples were measured. METABRIC dataset consists of images from three breast cancer grades, grade 1 (50 images), grade 2 (181 images) and grade 3 (269 images). Ali *et al*. segmented the single cells in the images using random forest classifier and then the expression of proteins in single cells was quantified. The mean protein expression of the segmented cells are used as the node features of the spatial graph with edge weights between cells defined using decaying kernels at the center of the reference cells. We used the following channels: HH3_total, CK19, CK8_18, Twist, CD68, CK14, SMA, Vimentin, c_Myc, HER2, CD3, HH3_ph, Erk1_2, Slug, ER, PR, p53, CD44, EpCAM, CD45, GATA3, CD20, Beta_catenin, CAIX, E_cadherin, Ki67, EGFR, pS6, Sox9, vWF_CD31, pmTOR, CK7, panCK, c_PARP_c_Casp3, DNA1, DNA2, H3K27me3, CK5, Fibronectin. Ali *et al*. annotated the following cell types: B cells, Basal CKlow, Endothelial, Fibroblasts, Fibroblasts CD68+, HER2+, HR+ CK7-, HR+ CK7-Ki67+, HR+ CK7-Slug+, HR-CK7+, HR-CK7-, HR-CKlow CK5+, HR-Ki67+, HRlow CKlow, Hypoxia, Macrophages Vim+ CD45low, Macrophages Vim+ Slug+, Macrophages Vim+ Slug-, Myoepithelial, Myofibroblasts and T cells, Vascular SMA+. We coarsened the cell types into B cells, Endothelial, Fibroblasts (Fibroblasts, Fibroblasts CD68+), Macrophages (Macrophages Vim+ CD45low, Macrophages Vim+ Slug+, Macrophages Vim+ Slug-), Myoepithelial, Myofibroblasts, T cells, Vascular SMA+ and Tumor cells (HER2+, HR+ CK7-, HR+ CK7-Ki67+, HR+ CK7-Slug+, HR-CK7+, HR-CK7-, HR-CKlow CK5+, HR-Ki67+, HRlow CKlow, Hypoxia).

### CODEX - colorectal cancer

The colorectal cancer dataset (Schürch *et al*.^7^ with 140 images from 35 patients) was measured with CODEX. The dataset consists of two patient groups, one group with Crohn’s-like reaction (CLR) represented in 68 images and one group with diffuse inflammatory infiltration (DII) represented in 72 images. Here, 57 proteins specific to the tumor microenvironment were measured. We used the segmentation previously performed by Schürch *et al*.. The molecular abundance per cell segment and the coordinates of the center of each cell were used to construct the spatial graph. We used the following channels: CD44, FOXP3, CD8A, TP53, GATA3, PTPRC, TBX21, CTNNB1, HLA-DR, CD274, MKI67, PTPRC, CD4, CR2, MUC1, TNFRSF8, CD2, VIM, MS4A1, LAG3, ATP1A1, CD5, IDO1, KRT1, ITGAM, NCAM1, ACTA1, BCL2, IL2RA, ITGAX, PDCD1, GZMB, EGFR, VISTA, FUT4, ICOS, SYP, GFAP, CD7, CD247, CHGA, CD163, PTPRC, CD68, PECAM1, PDPN, CD34, CD38, SDC1, HOECHST1:Cyc_1_ch_1, CDX2, COL6A1, CCR4, MMP9, TFRC, B3GAT1, MMP12. Schürch *et al*. annotated the following cell types: B cells, CD11b+ monocytes, CD11b+CD68+ macrophages, CD11c+ DCs, CD163+ macrophages, CD3+ T cells, CD4+ T cells, CD4+ T cells CD45RO+, CD4+ T cells GATA3+, CD68+ macrophages, CD68+ macrophages GzmB+, CD68+CD163+ macrophages, CD8+ T cells, NK cells, Tregs, adipocytes, dirt, granulocytes, immune cells, immune cells / vasculature, lymphatics, nerves, plasma cells, smooth muscle, stroma, tumor cells, tumor cells / immune cells and undefined, vasculature. We binarized the cell types into *immune cells* (B cells, CD11b+ monocytes, CD11b+CD68+ macrophages, CD11c+ DCs, CD163+ macrophages, CD3+ T cells, CD4+ T cells, CD4+ T cells CD45RO+, CD4+ T cells GATA3+, CD68+ macrophages, CD68+ macrophages GzmB+, CD68+CD163+ macrophages, CD8+ T cells, NK cells, Tregs, granulocytes, immune cells, immune cells / vasculature, lymphatics and tumor cells / immune cells) and *other* (adipocytes, dirt, nerves, plasma cells, smooth muscle, stroma, tumor cells, undefined and vasculature).

### Processing

We considered molecular feature spaces and cell type feature spaces. We compared molecular feature quantification per cell as provided by the authors^9,16^ (no feature transformation), image-wise feature standardization and global feature standardization. The cell type feature space was assembled as a one-hot encoding of the categorical cell type labels as described in the methods sections specific to each dataset. We used patient identifiers as the domain label for all datasets.

### Spatial proximity graphs

We considered neighborhood graphs built with fixed kernel radii across all images. In all datasets considered here, pixel dimensions are fixed across images so that radii defined on pixels correspond to consistent spatial distances across images. We defined a raw adjacency matrix *A* for each image with entries *a*_*ij*_ based on a radius *r* of a kernel between the position of two cells *i, j* in 2D space *z*_*i*_, *z*_*j*_

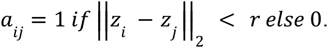

### Spectral clustering

We computed spectral clustering for the spatial graphs, where we divided the graph into a certain number of subgraphs (clusters) based on distances between the nodes of the graph. We first compute *k* nearest neighbors matrices, where *k* is a hyperparameter representing the number of neighbors used for kNN graph construction. We then computed the *n* spectral clusters in the graph using the *SpectralClustering* model, where *n* is a hyperparameter of the desired number of subgraphs and we perform a one-hot encoded assignment of the graph nodes to the nearest cluster. We also calculate the adjacency matrices within each of the clusters.

**Table 1:** Hyper-parameters screened in grid search for each data set.

| dataset | learning rate | l2 | radius | aggregation | depth | number of clusters | width |
| --- | --- | --- | --- | --- | --- | --- | --- |
| **IMC - breast cancer (Jackson)** | {**0.05**, 5e-3, 5e-4} | {**0**, 1e-6, 1e-3} | {**10**,50} | mean | {**1,2**,3} | {**5,10**, 20} | {**16**,64} |
| **IMC - breast cancer (METABRIC)** | {**5e-3**, 5e-4} | {**0**, 1e-6} | {10, **20**, 55} | mean | 3 | {**5**, 10} | {16, **64**} |
| **CODEX - colorectal cancer** | {**5e-3**, 5e-4} | {**0**, 1e-6} | {25,**50**, 120} | spectral | {**2,3**} | {**5**,10} | {4,8,**16**, 32} |

## Models

All models presented are feed-forward networks that take graph data (or reductions thereof) as input and produce a graph-level classification as output.

### Pseudobulk multi-layer perceptron networks (MLP)

For the pseudobulk reference model, we used an aggregated mean of the feature space across the entire image and it as input, 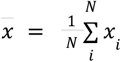, where 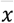 is the vector of average expression of input features, *x* is the node feature matrix (number of nodes *N x* input features). The input is then fed to a dense fully connected network to obtain the graph-level prediction 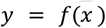.

### Multi-instance networks (MI)

For the multi-instance reference model, we used a stack of fully connected layers to obtain node-wise feature embeddings. Each layer *l* transformed the set of node features as:

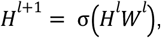

where *σ* is an activation function, *H*^*l*^ is a node feature matrix of dimensions (number of nodes x input features) and *W*^*l*^ is a weight matrix of dimensions (input features x output features). To then obtain graph-level predictions, the node feature embeddings were aggregated by a pooling layer and further transformed by two dense layers.

### Graph convolutional networks (GCN)

The node embedding layers for the Graph Convolutional Network are defined as:

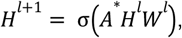

where *σ* is an activation function, *H*^*l*^ is a n input node feature matrix of dimensions (number of nodes x input features), *W*^*l*^ is a weight matrix of dimensions (input features x output features) and *A*^*^ is the spectral normalized adjacency matrix:

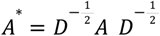

where *A* is the raw adjacency matrix and D is the degree matrix of A. Node embeddings were then aggregated by a pooling layer and two dense layers were then used to obtain the graph-level predictions.

### Graph convolutional networks with self-supervision (GCN-SS)

We introduced a self-supervision auxiliary task to the graph neural network. The self-supervision loss was added to the supervised loss which in this case acts as a regularization factor to the loss function. Here, we chose relative cell type proportions as our self-supervision task. Spectral clustering was performed on the spatial graph. For each node, the relative proportions of cell types in the same cluster are predicted.

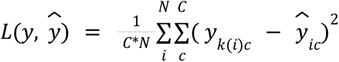

Where *y* ∈ *R*^*KxC*^, *ŷ* ∈ *R*^*NxC*^, where *K* is the number of spectral clusters and *k*(*i*) the cluster assignment of cell *i*, and *C* is the number of distinct cell types.

### Dispersion model

As for deriving self-supervision labels, we performed spectral clustering, calculated the relative proportions of cell types per cluster, and finally aggregated the mean of relative proportions of cell types across the clusters, 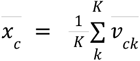, a vector of length *C*, where *K* is the number of spectral clusters and 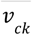 is the fraction of cell types of type *c* in cluster *k*. We then predicted the graph-level label based on 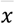 with a fully connected network: 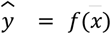. In this manuscript, we used the dispersion model only on binarized cell type labels.

### Node-wise pooling

We aggregated information across all observations in an image using a mean aggregation for the IMC datasets and used aggregation in spectral clusters on the CODEX data. For spectral pooling, we first aggregated nodes within the same cluster followed by aggregating across clusters. The spectral clusters are subgraphs (clusters) of adjacent nodes assigned k-nearest neighbors clustering algorithm. The number of clusters in a graph is a hyperparameter ranging between 5 and 20 clusters.

### Integration metrics

To quantify the domain correction across the different models, we adapted integration metrics from scRNA-seq, namely the *graph connectivity* metric and *iLISI graph* metric from the scIB^9^ package. The *graph connectivity* and *iLISI graph* metrics are both calculated from the kNN graph (k=50) of the node embeddings of all images and assess the connectivity between the nodes from the same images, the higher the scores the better the images are integrated. We further trained a linear regression model using the node embeddings from the last layer of the models for the prediction of the image identities and used its accuracy to quantify the strength of the signal still coming from the image identity. For the *prediction accuracy*, the lower the score the better the images are integrated.

## Code and data availability

We used published datasets provided in the original studies. We summarized all models, training and interpretation mechanisms discussed here in a python package centered around graph-level supervision on spatial single-cell graphs, https://github.com/theislab/tissue. We are also providing all the analysis and the model evaluation notebooks that were presented throughout the paper, https://github.com/theislab/tissue_reproducibility.

## Author contributions

DSF and FJT conceived the study. DSF, MA and SR implemented the overall software and performed the analyses. DSF, MA, SR, AE and FJT wrote the manuscript.

## Acknowledgements

We would like to thank Jana R. Fischer, Jonas Windhager, and Prof. Dr. Bernd Bodenmiller for assistance with the discussed breast cancer data sets and for discussion on the topic of graph convolutional networks and cancer grade prediction. We would like to thank Eeshit Dhaval Vaishnav Prof. Dr. Aviv Regev for discussion on spatial molecular profiling data. We would like to thank Anna Schaar for discussions about the model and the code base.

This work was supported by the German Federal Ministry of Education and Research (BMBF) under Grant No. 01IS18036B and No. 01IS18053A, by the Wellcome Trust Grant 108413/A/15/D and by the Helmholtz Association’s Initiative and Networking Fund through Helmholtz AI [grant number: ZT-I-PF-5-01]. D.S.F. acknowledges support from a German Research Foundation (DFG) fellowship through the Graduate School of Quantitative Biosciences Munich (QBM) [GSC 1006 to D.S.F.] and by the Joachim Herz Foundation. S. R. is supported by the Helmholtz Association under the joint research school “Munich School for Data Science”— MUDS.

## Conflicts of interest

F.J.T. consults for Immunai Inc., Singularity Bio B.V., CytoReason Ltd, and Omniscope Ltd, and has ownership interest in Dermagnostix GmbH and Cellarity.

