## Supplementary Figure 1-9 for "Graph neural networks learn emergent tissue properties from spatial molecular profiles"

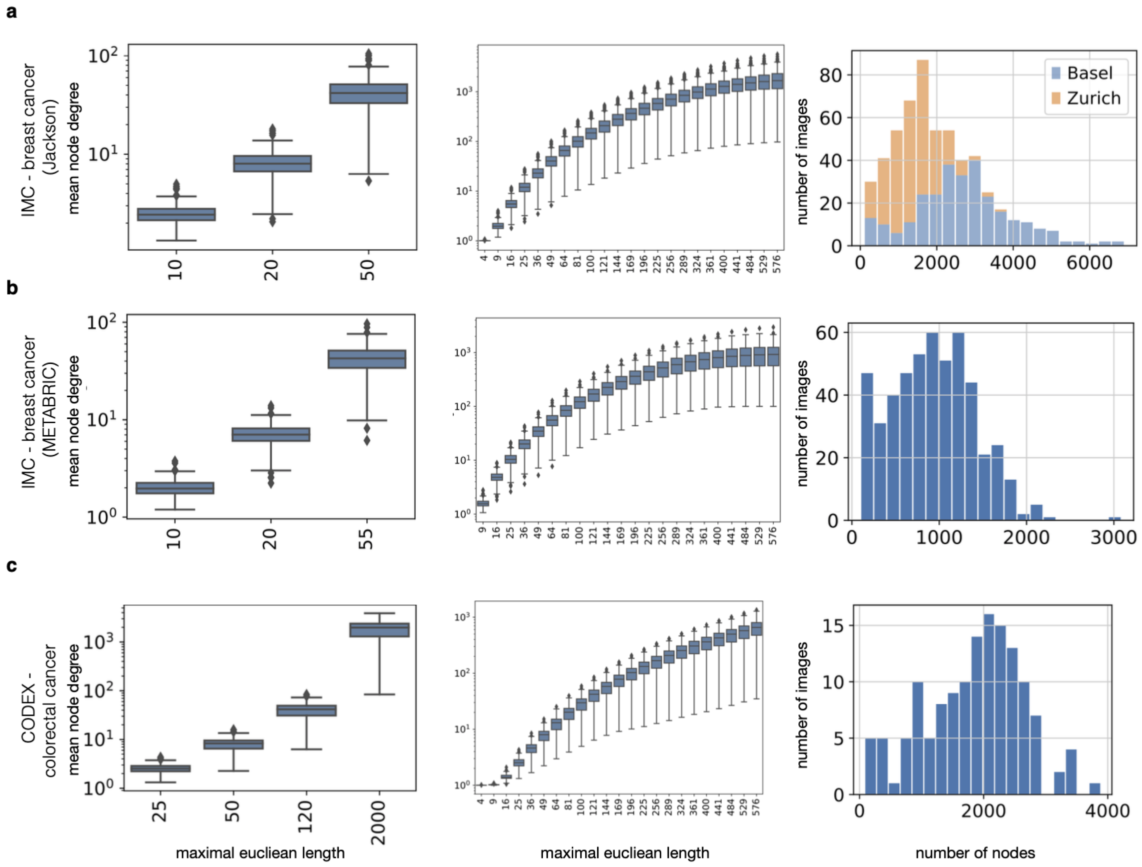


**Supp. Fig. 1: Graph summary statistics of analyzed data.**

Mean node-degree per image by radius length used in benchmarks (left) and in a scan across (middle), and number of nodes per image (right) for **(a)** IMC - breast cancer (Jackson), **(b)** IMC - breast cancer (METABRIC) and **(c)** CODEX - colorectal cancer datasets.

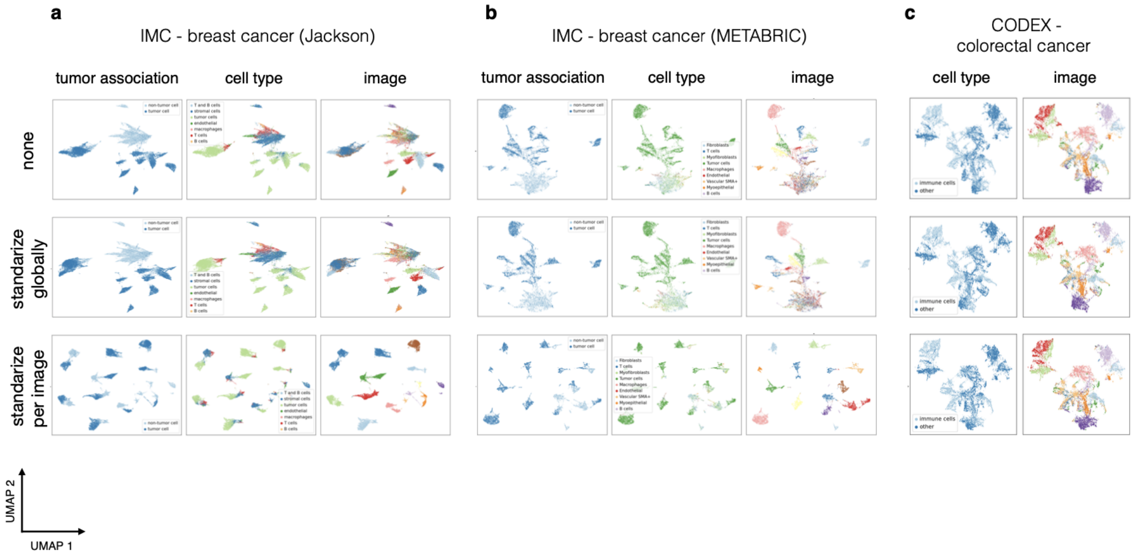


**Supp. Fig. 2: Node preprocessing.**

Coarse cell type group, cell types and image identity superimposed on UMAP learned on molecular features after different feature processing for subsampled **(a)** IMC - breast cancer (Jackson), **(b)** IMC - breast cancer (METABRIC) and **(c)** CODEX - colorectal cancer datasets.

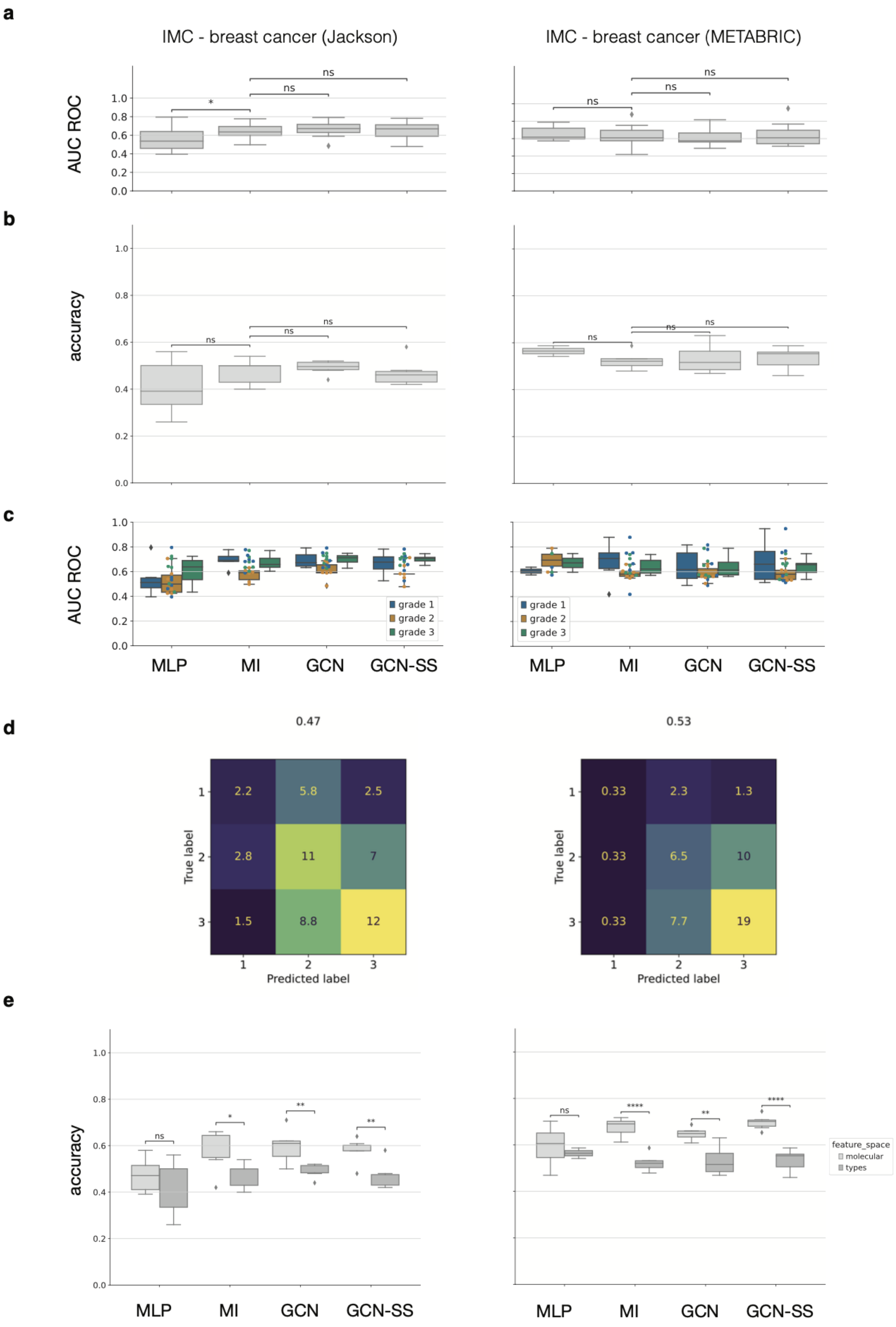
a
IMC - breast cancer (Jackson)
IMC - breast cancer (METABRIC)
AUC ROC
ns
*
b
accuracy
c
grade 1
grade 2
grade 3
MLP
MI
GCN
GCN-SS
d
0.47
0.53
True label
Predicted label
2.2
5.8
2.5
2.8
11
7
1.5
8.8
12
0.33
2.3
1.3
0.33
6.5
10
0.33
7.7
19
e
feature_space
molecular
types
**
****

**Supp. Fig. 3: Predicting tumor phenotypes from categorical cell type representations of nodes on breast cancer datasets.**

Shown are three separate applications of graph neural networks to predict tumor phenotypes on the IMC - breast cancer (Jackson) and IMC - breast cancer (METABRIC). **(a, b)** Model complexity ablation study on classification performance on breast cancer grade prediction. Shown is **(a)** area-under-curve of the receiver-operator characteristic curve (AUC ROC) for categorical cell types, **(b)** the accuracy and **(c)** area-under-curve of the receiver-operator characteristic curve (AUC ROC) for each class across six-fold cross-validation for the best performing hyper-parameter set for each model class. *MLP*: Multi-layer perceptron based on graph-wide summary statistics on features. *MI*: Multi-instance model on nodes of graph. *GCN*: Graph convolutional network. *GCN-SS*: Graph convolutional network with additional self-supervision loss. **(d)** Mean confusion matrix of best performing network across cross validation partitions for the best performing GCN-SS. **(e)** Comparison between the performance of the molecular and types features for the different datasets.

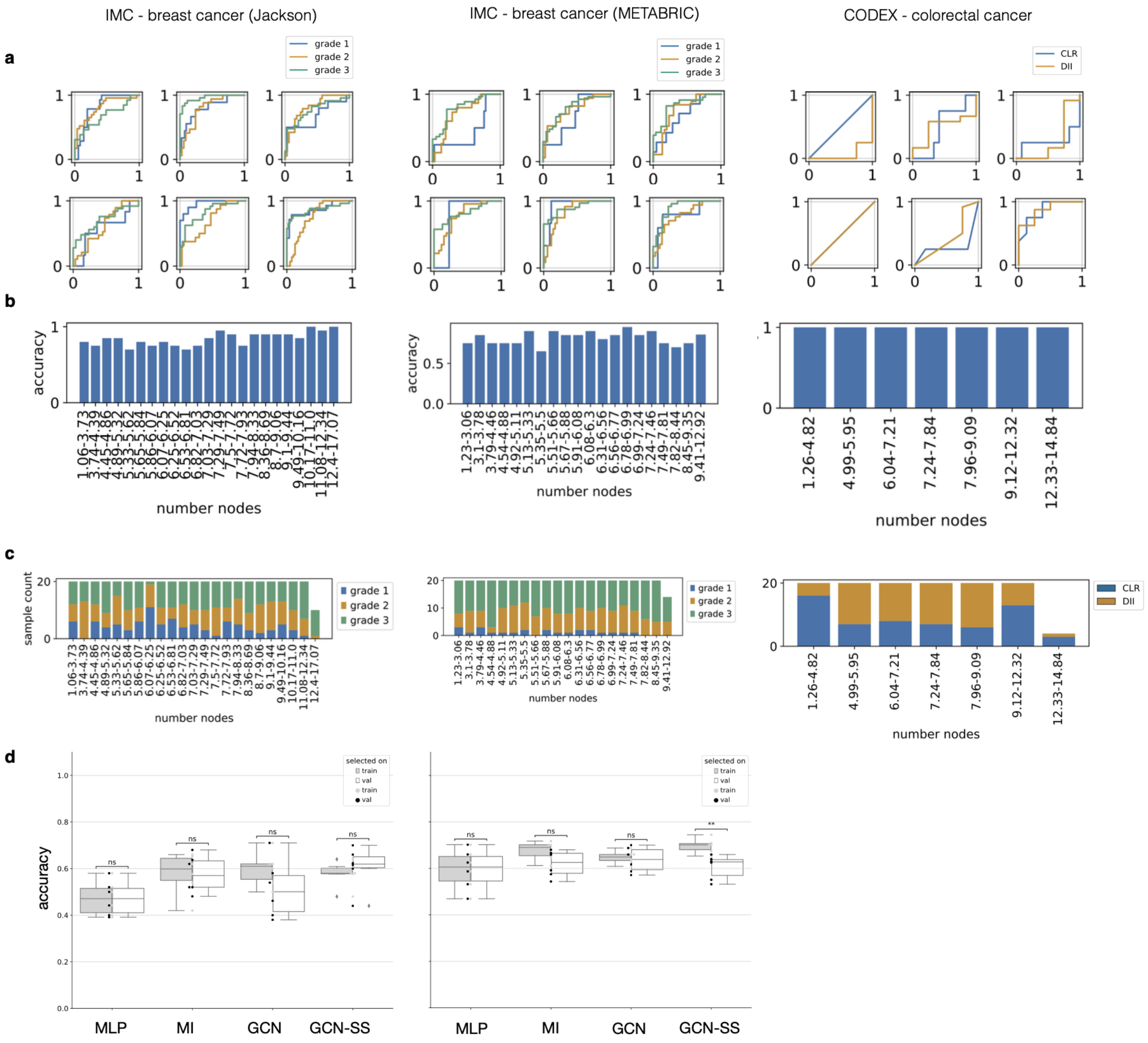


**Supp. Fig. 4: Detailed performance analysis of models based on a molecular feature space.**

**(a)** ROC curves for prediction task for the different folds of cross validation, **(b)** binned accuracy by number of nodes in graph and **(c)** label distribution over graph size bins for IMC - breast cancer (Jackson), IMC - breast cancer (METABRIC) and CODEX - colorectal cancer datasets. **(d)** Comparison between the accuracy of the different models when selected on train vs validation losses.

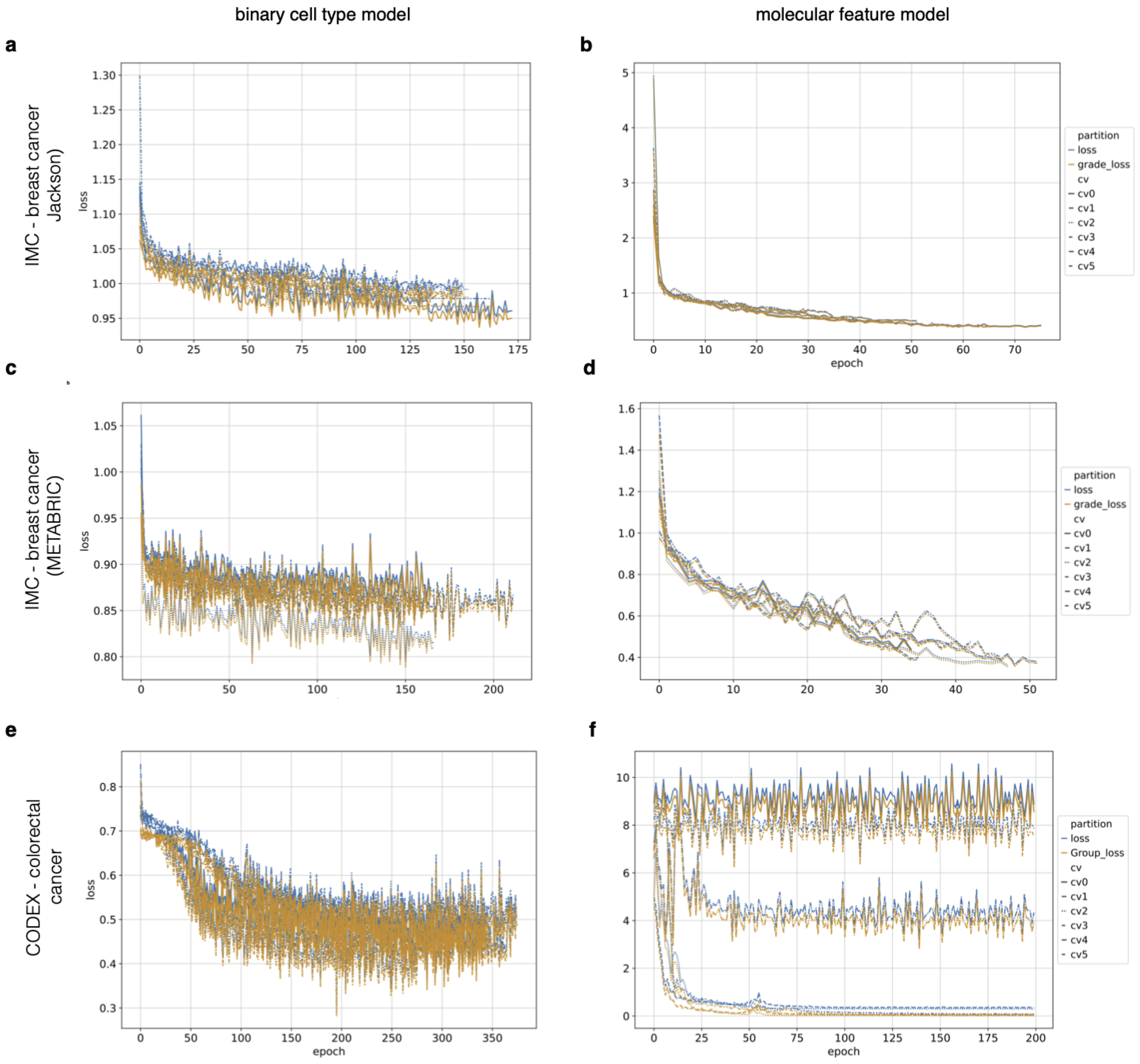


**Supp. Fig. 5: Training of models that predict cancer grade based on molecular feature space.**

Epoch-wise training plots for best performing binary cell types and molecular feature space-based model of GCN-SS model class selected based on mean cross-entropy loss on train data across cross-validations: for the different datasets: **(a, b)** IMC - breast cancer (Jackson), **(c, d)** IMC - breast cancer (METABRIC) and **(e, f)** CODEX - colorectal cancer.

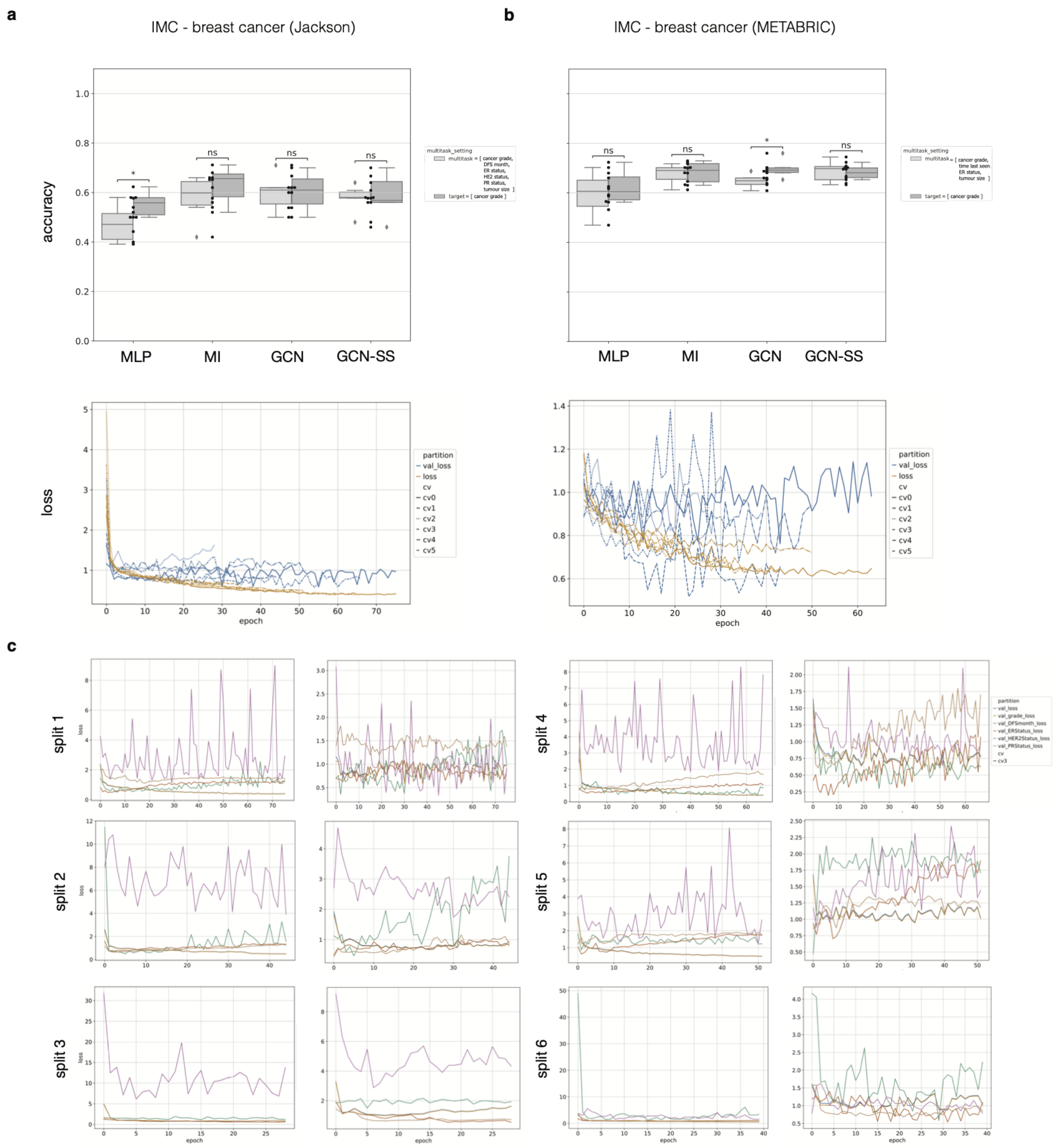


**Supp. Fig. 6: Multitasking models.**

On breast cancer datasets, **(a, b)** shown is the accuracy on test set based on number of tasks (upper panel) and epoch-wise training of multi-tasking models (lower panel) on **(a)** IMC - breast cancer (Jackson) dataset, and **(b)** IMC - breast cancer (METABRIC). *target* with only the cancer grade and *multitask* with cancer grade, disease-free survival (DFS) month, estrogen receptor (ER) status HER2 status, progesterone receptor (PR) status and tumor size. **(c)** Epoch-wise training plot for each task in multi-tasked setting for both training and validation partition for 6 cross validations on IMC - breast cancer (Jackson).

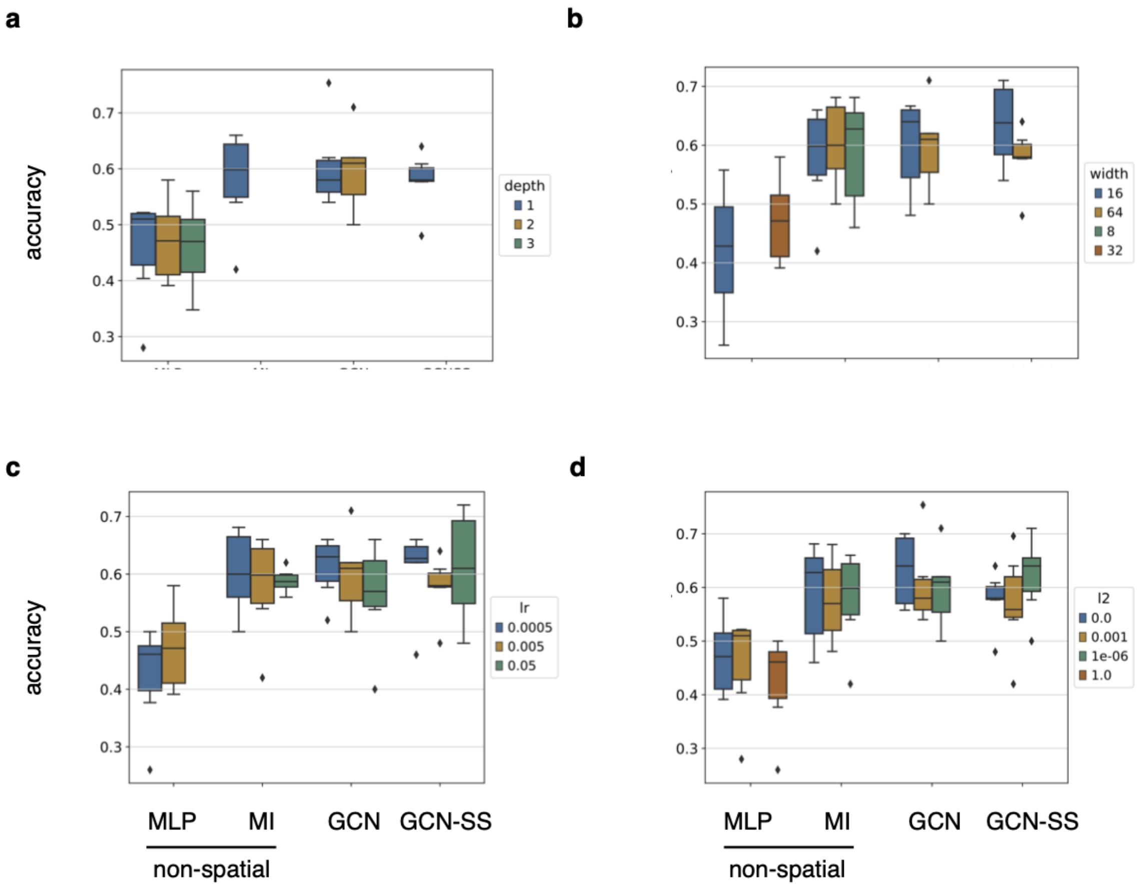


**Supp. Fig. 7: Model topology comparison.**

On IMC - breast cancer (Jackson), we plot best models test accuracy by hyper parameter partition with x axis as the different models, MLP, MI, GCN and GCN-SS: **(a)** depth, **(b)** width of dense layer, **(c)** learning rate and **(d)** L2 regularization factor. *MLP*: Multi-layer perceptron based on graph-wide summary statistics on features. *MI*: Multi-instance model on nodes of graph. *GCN*: Graph convolutional network. *GCN-SS*: Graph convolutional network with additional self-supervision loss.

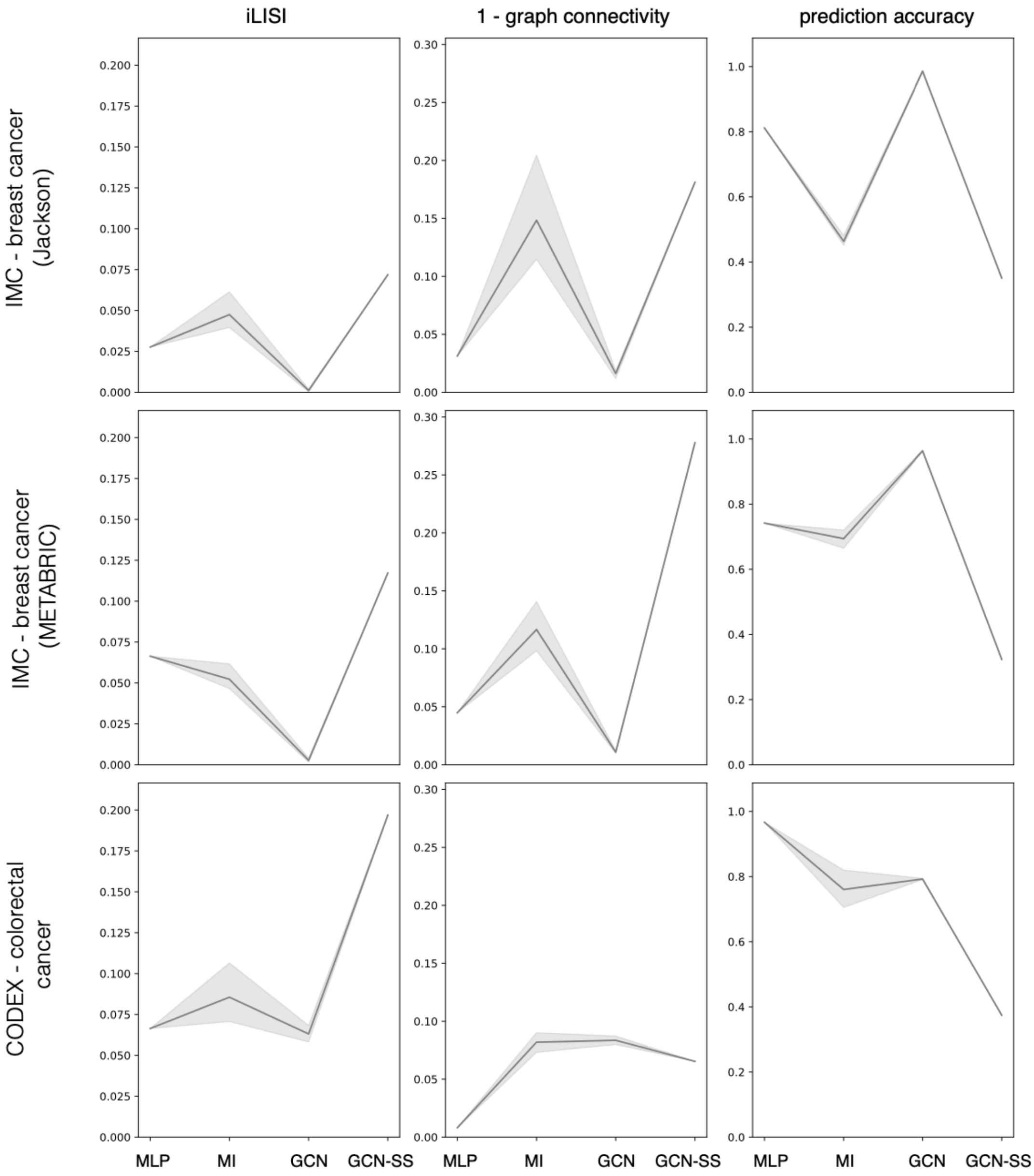


**Supp. Fig. 8: integration metrics**

Shown are metrics measuring the domain correction on the different datasets IMC - breast cancer (Jackson), IMC - breast cancer (METABRIC) and CODEX - colorectal cancer using different measures (N=3 cross validations per method and dataset). *iLISI graph*, *graph connectivity* and *regression prediction accuracy,* across three cross-validations.

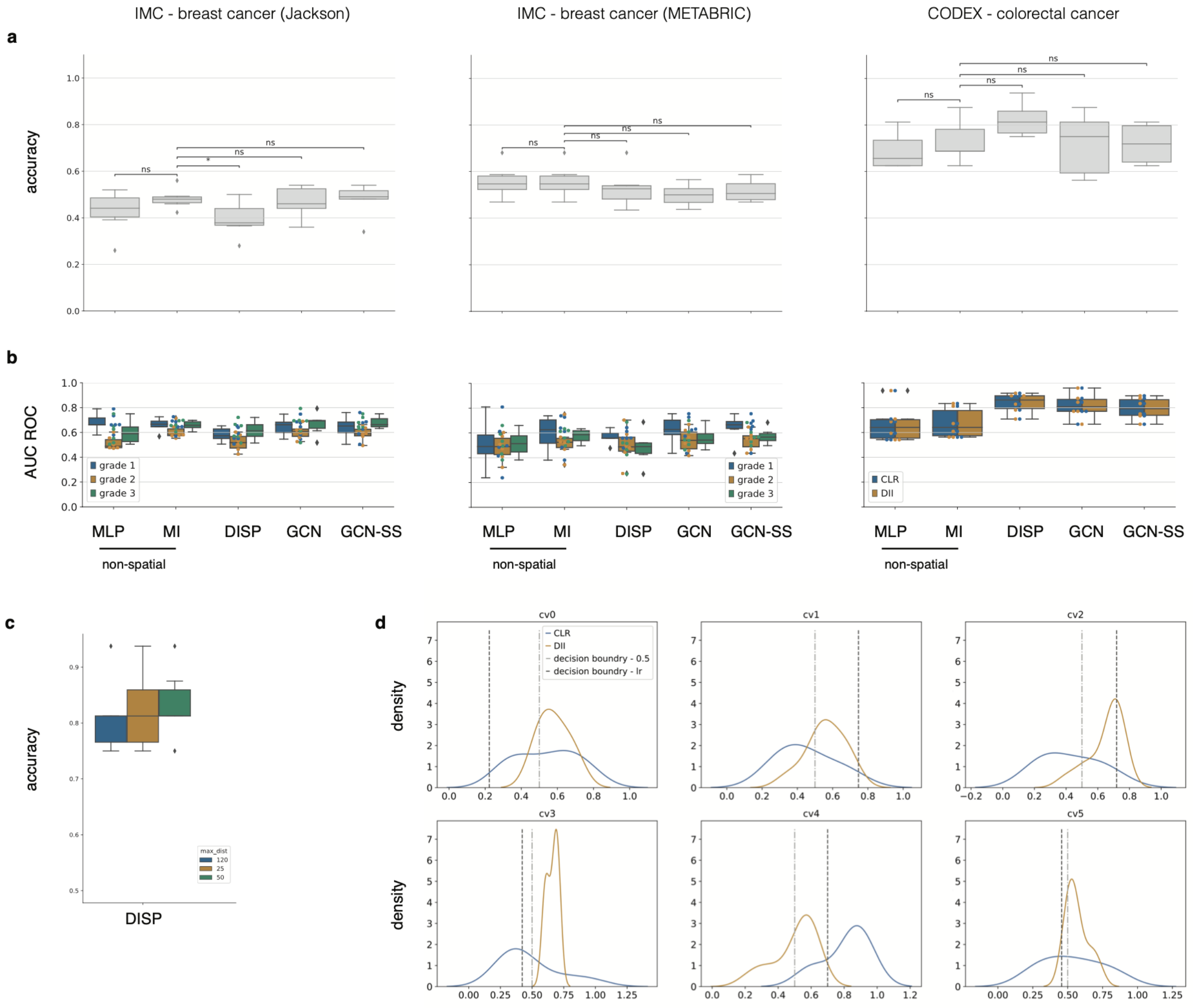


**Supp. Fig. 9: Performance of tumor classification using binary cell-type feature space**

Shown are three separate applications to predict tumor phenotypes on the IMC - breast cancer (Jackson), IMC - breast cancer (METABRIC) and CODEX - colorectal cancer datasets using binary cell type feature space **(a, b)**. Shown are **(a)** the accuracy to predict tumor phenotypes across different models and **(b)** the area-under-curve of the receiver-operator characteristic curve for each tumor grade across six-fold cross-validation for the best performing hyper-parameter set for each model class for best models selected on train loss. **(c, d)** On on the CODEX - colorectal cancer dataset, we show **(c)** the accuracy of the different niche radii of dispersion model (DISP) and **(d)** the distribution of the two tumor classes, Crohn's-like reaction (CLR) and diffuse inflammatory infiltration (DII) highlighting the 0.5 decision boundary and the logistic regression decision boundary.
